# FERONIA-Dependent Translational Buffering of Ribosome-Associated Genes in Salt-Stressed Tomato Roots

**DOI:** 10.64898/2026.06.11.731639

**Authors:** Junyu Bai, Yanfen Fan, Ruolin Yang, Jingquan Yu

## Abstract

Salt stress limits tomato productivity, yet how translational regulation contributes to root salt adaptation remains poorly understood. We integrated RNA-seq and ribosome profiling in wild-type (WT) and *fer* tomato roots under control and 150 mM NaCl conditions. In WT roots, the salt response was predominantly transcript-driven, but a 29-gene ribosome-associated module showed reduced RNA abundance alongside increased translational efficiency (TE), indicating selective translational buffering. FER deficiency disrupted this balance: it constitutively elevated ribosome occupancy of ribosome-associated genes while reducing basal expression of stress- and ion-transport-related genes. Under salt treatment, *fer* further showed stronger ion-transport transcriptional responses but weaker ribosome-associated TE responses, consistent with impaired translational selectivity. WT salt stress also shifted ribosome allocation away from the 5^′^UTR toward the CDS, an effect attenuated in *fer*; upstream open reading frame (uORF) and coding sequence (CDS) translational efficiency were also positively coupled during salt treatment. Feature modelling confirmed established features associated with uORF translation and further identified weaker local RNA folding around the uORF start codon, proline and charged amino acid enrichment, and TAG or TGA stop codon identity as positive predictors. Together, these results reveal FER-dependent changes in ribosome-associated translational buffering during tomato root salt responses.

## 1 Introduction

Salt stress is one of the most serious abiotic constraints on tomato production because cultivated tomato is generally moderately salt-sensitive: excess NaCl suppresses biomass accumulation, fruit set and yield, and reshapes fruit-quality traits [1, 2]. Our previous work in tomato has shown that RBOH1-dependent H_2_O_2_ production contributes to salt-stress acclimation and cross-tolerance, highlighting ROS signalling as an established component of tomato salt responses [3, 4]. In this context, roots are especially important because they are the first organ to encounter osmotic and ionic stress and thus determine Na^+^ exclusion, K^+^ retention, hydraulic adjustment and root-to-shoot signalling [5, 6]. In tomato roots, salinity reshapes root system architecture and induces rapid transcriptome and alternative-splicing reprogramming, supporting roots as an informative tissue for early stress-response analysis [7, 8].

Plant stress studies have historically relied heavily on transcriptome analyses to define stress-responsive genes and pathways, and recent work has increasingly expanded toward multi-omics integration of genomic, proteomic, metabolomic, epigenomic and phenomic data [9–12]. However, mRNA abundance often correlates imperfectly with protein output, indicating that post-transcriptional control, particularly translational regulation, can substantially reshape plant stress responses. Ribosome profiling (Ribo-seq) has therefore become a key approach because it captures in planta ribosome-protected footprints and, when paired with RNA-seq, allows quantification of ribosome occupancy and translational efficiency (TE) [13, 14]. Translatome-level studies of stress responses remain at an early stage, with examples including rice salinity, potato drought/heat and lychee cold stress [15–18]. A tomato root translatome has been mapped under non-stress conditions, demonstrating that Ribo-seq can refine annotation and reveal widespread non-canonical translation in this crop; what remains largely unknown is how salt stress rewires translation in tomato roots [19].

FERONIA (FER) is a conserved member of the *Catharanthus roseus* receptor-like kinase 1-like (CrRLK1L) family and functions at the cell surface as a multifunctional signalling hub [20]. FER was initially linked to growth control and root hair development through ROP/RAC signalling [21], and was later shown to mediate RALF-dependent root growth inhibition [22] and immune-complex organisation [23]. In tomato roots, the RALF2–FER–MYB63 module further connects FER signalling with growth–defence coordination [24], supporting its role as a receptor that integrates growth, defence and environmental information rather than a pathway-specific factor. Current reviews and integrated-omics analyses place FER at the centre of an extensive signalling network connecting cell-wall status, peptide ligands, hormone crosstalk, Ca^2+^ and ROS signalling, and developmental plasticity [20, 25]. In salinity responses, FER maintains cell-wall integrity during salt stress through a Ca^2+^-dependent mechanism and promotes salt tolerance by controlling photorespiratory flux [26, 27]. FER has also been connected to translation-related pathways: RALF1–FER phosphorylates eIF4E1 to promote mRNA translation and protein synthesis in Arabidopsis root hairs, and RALF1–FER interacts with and activates TOR signalling under low-nutrient conditions [28, 29].

A relevant layer for examining translational control is provided by 5^′^ leaders and upstream open reading frames (uORFs). uORFs are widespread elements in plant 5^′^ leaders that can conditionally regulate translation of the downstream main ORF by altering scanning, reinitiation or ribosome stalling [30]. In plants, uORFs are not merely passive repressors: the TBF1 leader links translation to the growth-to-defence transition, and SAC51/SACL1 transcripts are translationally regulated in response to thermospermine through uORF-dependent control [31, 32]. More broadly, conserved peptide uORFs can mediate stress-responsive translational control across plant lineages [33]. Their regulatory effects are shaped by start-codon context, uORF arrangement and spacing, leader architecture, and natural sequence variation [34, 35]. Ribo-seq data, when combined with ORF-calling approaches such as RiboCode, PRICE and RiboBA [36–38], enable translated uORFs to be identified in plant transcriptomes [19, 39, 40].

These observations led us to ask whether FER participates in salt-responsive translational regulation in tomato roots. We treated wild-type and *fer* seedlings under control conditions or 150 mM NaCl treatment and sequenced matched RNA-seq and Ribo-seq libraries from root tissues. In WT roots, salt-stress regulation was largely transcript-driven but showed incomplete transcription–translation coupling, with ribosome/translation-related genes forming a prominent translationally buffered class. FER deficiency increased basal transcript abundance and ribosome occupancy of ribosome/translation-related genes, weakened their salt-induced TE responsiveness, and altered FER-dependent ion-transport responses. We also identified features favouring translated uORFs, including shorter relative distance from the CDS, higher uORF GC content, higher proline and charged amino-acid proportions, and region-specific RNAfold-derived folding metrics. These results provide a catalogue of non-canonical ORF translation events in tomato and reveal a role for FER in salt-responsive translational regulation in tomato roots.

## 2 Results

### 2.1 Salt Stress Phenotypes and Quality Assessment of RNA-seq and Ribo-seq Libraries

To examine transcriptomic and translatomic responses associated with FER-dependent salt stress regulation, we performed RNA-seq and ribosome profiling using roots of WT and *fer* seedlings grown under control or 150 mM NaCl conditions. The four experimental groups were WT-CK (WC), WT-S150 (WS), *fer*-CK (FC), and *fer*-S150 (FS), each with three biological replicates. Under control conditions, *fer* seedlings already showed reduced fresh weight compared with WT, indicating a basal growth defect (Figure 1A–B). Salt treatment further inhibited growth in both genotypes; however, the relative reduction in fresh weight was significantly greater in *fer* than in WT (Figure 1B), indicating enhanced salt sensitivity of the mutant.

**Figure 1:**
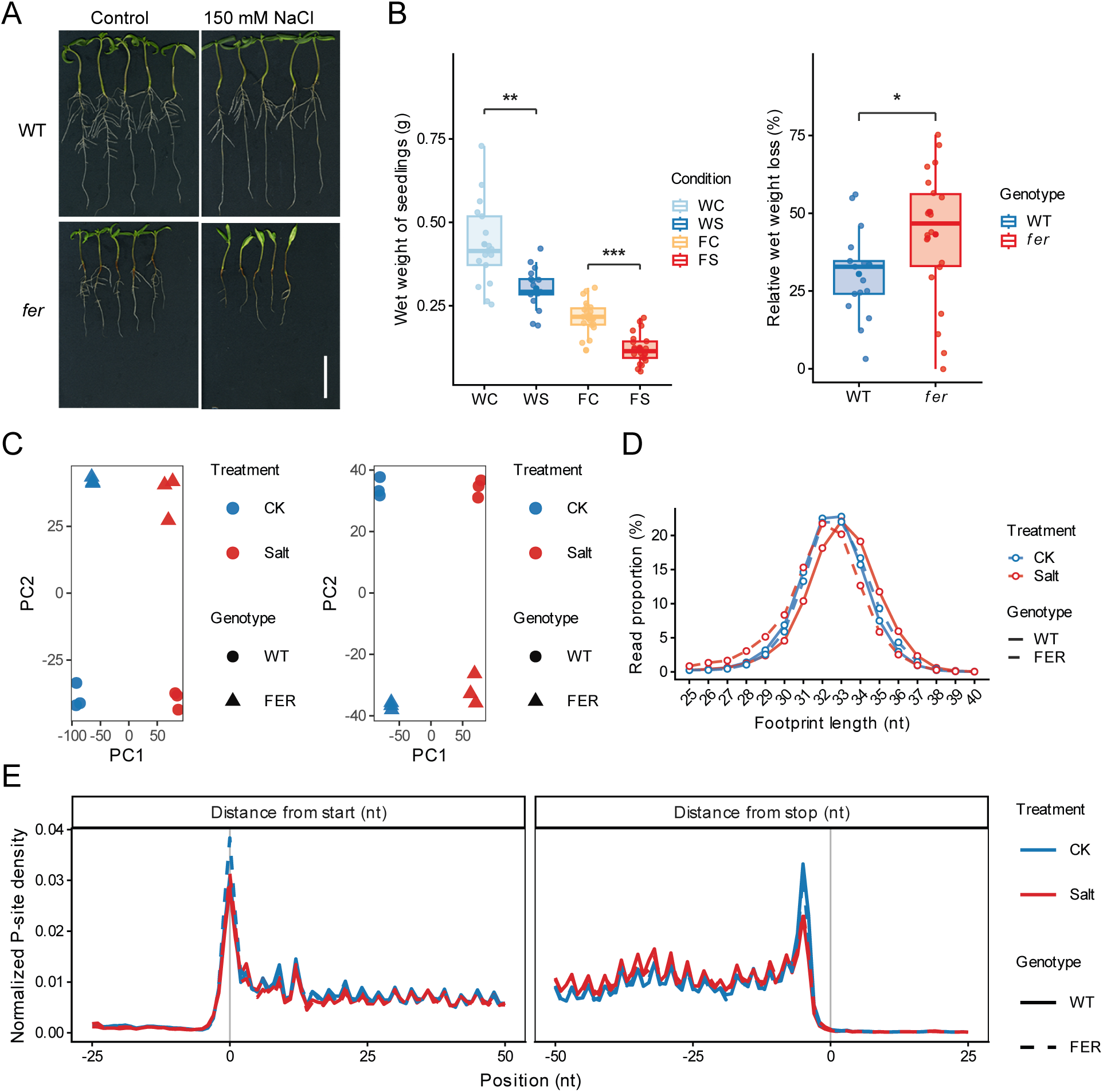
Phenotypes and sequencing quality of WT and *fer* tomato seedlings under control and salt-stress conditions. **(A)** WT and *fer* seedlings grown under control conditions or 150 mM NaCl treatment. **(B)** Seedling wet weight and relative wet weight change under salt stress across WC, WS, FC and FS conditions. **(C)** Principal component analysis (PCA) of RNA-seq and Ribo-seq libraries. **(D)** Length distribution of ribosome-protected fragments across the four experimental groups. **(E)** Metagene profiles of normalized P-site density around start and stop codons. Asterisks indicate significance levels: * *P* < 0.05, ** *P <* 0.01, *** *P <* 0.001; two-sided Wilcoxon rank-sum tests.

In total, approximately 361 million, 402 million, 390 million, and 377 million ribosome profiling reads were obtained from WC, WS, FC, and FS samples, respectively, and approximately 81 million, 67 million, 70 million, and 65 million RNA-seq reads from the corresponding samples (Tables S1 and S2). After rRNA removal, 24.79%–47.43% of Ribo-seq reads mapped to the tomato reference genome (SL4.0/ITAG4.0). Pairwise Pearson correlation of transcript abundance and ribosome occupancy showed high within-condition reproducibility, with *R*^2^ *>* 0.95 for RNA-seq and *R*^2^ *>* 0.93 for Ribo-seq libraries across biological replicates (Supplementary Figure S1A). Principal component analysis further confirmed that replicates clustered closely within conditions and that samples separated clearly by treatment and genotype, with treatment effects more pronounced than genotype effects at both regulatory levels (Figure 1C).

Ribosome profiling data showed the expected characteristics of high-quality actively translated libraries. Ribosome-protected fragments (RPFs) were mainly enriched at 31–35 nt, with the dominant peak around 33 nt across all four groups (Figure 1D). Frame 0 was the dominant reading frame in all groups, accounting for approximately 38.9%–40.3% of CDS P-sites, indicating clear three-nucleotide periodicity (Supplementary Figure S1B). Metagene analysis showed a sharp P-site peak at annotated start codons and enrichment near stop codons, supporting reliable P-site calibration (Figure 1E). Ribo-seq P-sites were predominantly assigned to CDS regions (96.2%–96.8% across the four groups), whereas RNA-seq reads were more broadly distributed across 5^′^UTR, CDS, and 3^′^UTR regions (Supplementary Figure S1C). Together, these results confirmed the quality and reproducibility of both datasets for subsequent analyses.

### 2.2 Regulatory Classification of the WT Salt Response Reveals Translational Buffering of Ribosomal Genes

Salt stress triggered extensive reprogramming in WT roots at both regulatory levels. In the WS/WC comparison, 1,174 up-regulated and 1,183 down-regulated genes were identified at the transcript abundance level (adjusted *p <* 0.05, *|* log_2_ FC*| ≥* 1), alongside 1,238 up- and 1,120 down-regulated genes at the ribosome occupancy level (Supplementary Figure S2A; Table S3); 1,016 induced and 939 repressed genes were shared across both layers (72.8% and 68.8% of their respective unions), yet a substantial fraction was regulated exclusively at one level (Supplementary Figure S2B), indicating that the WT salt response involves layer-specific events beyond transcriptional control. To characterise how these changes are coordinated, we applied the deltaTE framework [14], classifying genes into four regulatory categories based on RNA, ribosome occupancy, and TE changes (Supplementary Figure S3A): Forwarded (ribosome occupancy tracks mRNA without a significant TE change), Buffered (TE changes oppose mRNA changes and attenuate ribosome occupancy changes), Intensified (TE changes reinforce mRNA changes), and Exclusive (significant TE and ribosome occupancy changes without a significant mRNA change). The Forwarded class was by far the most abundant in both directions (2,842 up and 2,894 down; Figure 2A), confirming that transcriptional reprogramming is the dominant driver of the WT salt response, while the Buffered (103 up, 276 down), Intensified (69 up, 44 down), and Exclusive (82 up, 73 down) classes defined subsets subject to additional translational regulation (inset, Figure 2A; Table S3).

**Figure 2:**
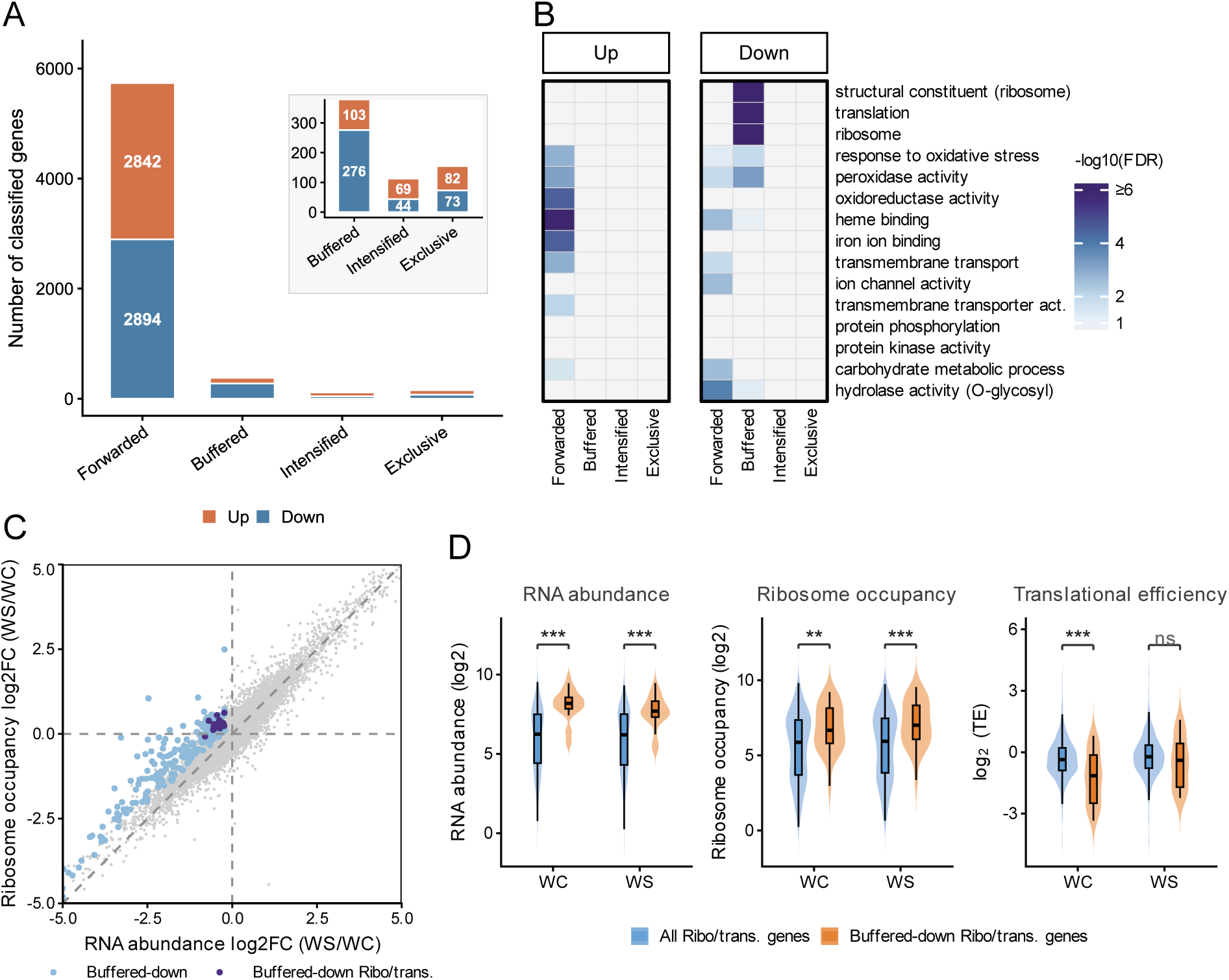
Transcription-translation concordance of salt-responsive genes in WT tomato roots. **(A)** Genes responsive in the WS/WC comparison classified into four regulatory classes. Up/down subsets for Forwarded, Buffered and Intensified classes are defined by RNA log_2_FC direction; for the Exclusive class, by Ribo log_2_FC direction. **(B)** Gene Ontology enrichment of genes in each regulatory class. **(C)** RNA and ribosome occupancy changes, highlighting buffered-down genes with and without ribosome/translation GO annotation. **(D)** RNA abundance, ribosome occupancy and translational efficiency of ribosome/translation-annotated genes and the buffered-down subset under WC and WS conditions. Asterisks in (D): ** *P <* 0.01, *** *P <* 0.001; two-sided Wilcoxon rank-sum tests.

GO enrichment analysis revealed functional signatures across regulatory classes (Figure 2B; Table S4). Forwarded-up genes were enriched for redox- and membrane transport-related functions, whereas Forwarded-down genes were enriched for carbohydrate metabolism, O-glycosyl hydrolase activity, and selected transport-related terms. The most prominent signal was observed in the Buffered-down class, where structural constituent of ribosome, translation, and ribosome were the highest-confidence terms (*−* log_10_ FDR *≥* 6), indicating that ribosome/translation-associated genes are concentrated among transcripts that are down-regulated at the RNA level but show opposing TE increases, a pattern expected to attenuate salt-induced decreases in ribosome occupancy.

To examine the gene-level behaviour underlying this enrichment, we highlighted Buffered-down genes in the genome-wide RNA versus ribosome occupancy log_2_FC scatter plot and further marked those annotated with ribosome/translation GO terms (Figure 2C; Table S6). Buffered-down genes occupied the region with negative RNA changes but comparatively modest ribosome-occupancy changes, consistent with TE increases attenuating the effect of transcript repression. The ribosome/translation-annotated subset followed this same buffered pattern but had a higher fraction of genes with ribosome occupancy increased in WS relative to WC than the broader Buffered-down set (28 of 29 versus 130 of 276), despite reduced RNA abundance in WS. Because the enriched Buffered-down genes represented only a subset of ribosome/translation-annotated genes, we next asked whether they differed from the broader ribosome/translation gene population in condition-level RNA abundance, ribosome occupancy, and TE under WC and WS conditions (Figure 2D). The Buffered-down subset showed higher RNA abundance and ribosome occupancy than the broader ribosome/translation gene set under both conditions, indicating that these genes constitute a highly expressed ribosome/translation submodule. Notably, this subset had significantly lower TE under WC, whereas its TE became comparable to that of the broader ribosome/translation gene set under WS. Together, these results indicate that these 29 ribosome/translation-annotated Buffered-down genes form a prominent WT salt-response module, in which salt-associated RNA reduction is accompanied by increased TE and, for most genes, increased ribosome occupancy.

### 2.3 Basal Transcription–Translation Differences in *fer* Roots Reveal FER-Dependent Regulation of Ribosomal Genes

Having defined the transcription–translation architecture of the WT salt response, including a Buffered-down ribosome/translation module, we next asked whether loss of FER had already altered this regulatory landscape before salt exposure. We therefore compared *fer* and WT roots under control conditions (FC/WC) to identify pre-existing FER-dependent differences in RNA abundance, ribosome occupancy and TE. Under control conditions, *fer* roots differed from WT at both regulatory layers: 384 up- and 293 down-regulated genes were detected at the transcript level, and 376 up-and 316 down-regulated genes at the ribosome occupancy level (Supplementary Figure S2C), with 67.4% and 51.1% of induced and repressed genes shared across both layers, respectively (Supplementary Figure S2D). Most classified genes belonged to the Forwarded class (*n* = 2,945), with minimal representation in the Buffered (*n* = 10), Intensified (*n* = 3), and Exclusive (*n* = 17) classes (Supplementary Figure S3B), indicating that FER deficiency reshapes basal gene expression primarily at the transcript level without widespread translational remodelling. GO enrichment analysis of the Forwarded-up class revealed a notable functional signal: structural constituent of ribosome, translation, and ribosome were the top enriched terms (*−* log_10_ FDR *≥* 6; Figure 3C). This indicates that ribosome/translation-associated genes are co-upregulated at both RNA and ribosome-occupancy levels in *fer* roots at baseline. The Forwarded-down class was enriched for heme binding, peroxidase activity, oxidative-stress response, and ion/transmembrane transport-related functions (Figure 3C), suggesting reduced basal expression of stress- and transport-associated genes in *fer*. This basal reduction is consistent with a weakened stress-preparedness state in *fer* roots, which may contribute to their enhanced sensitivity to subsequent salt exposure.

**Figure 3:**
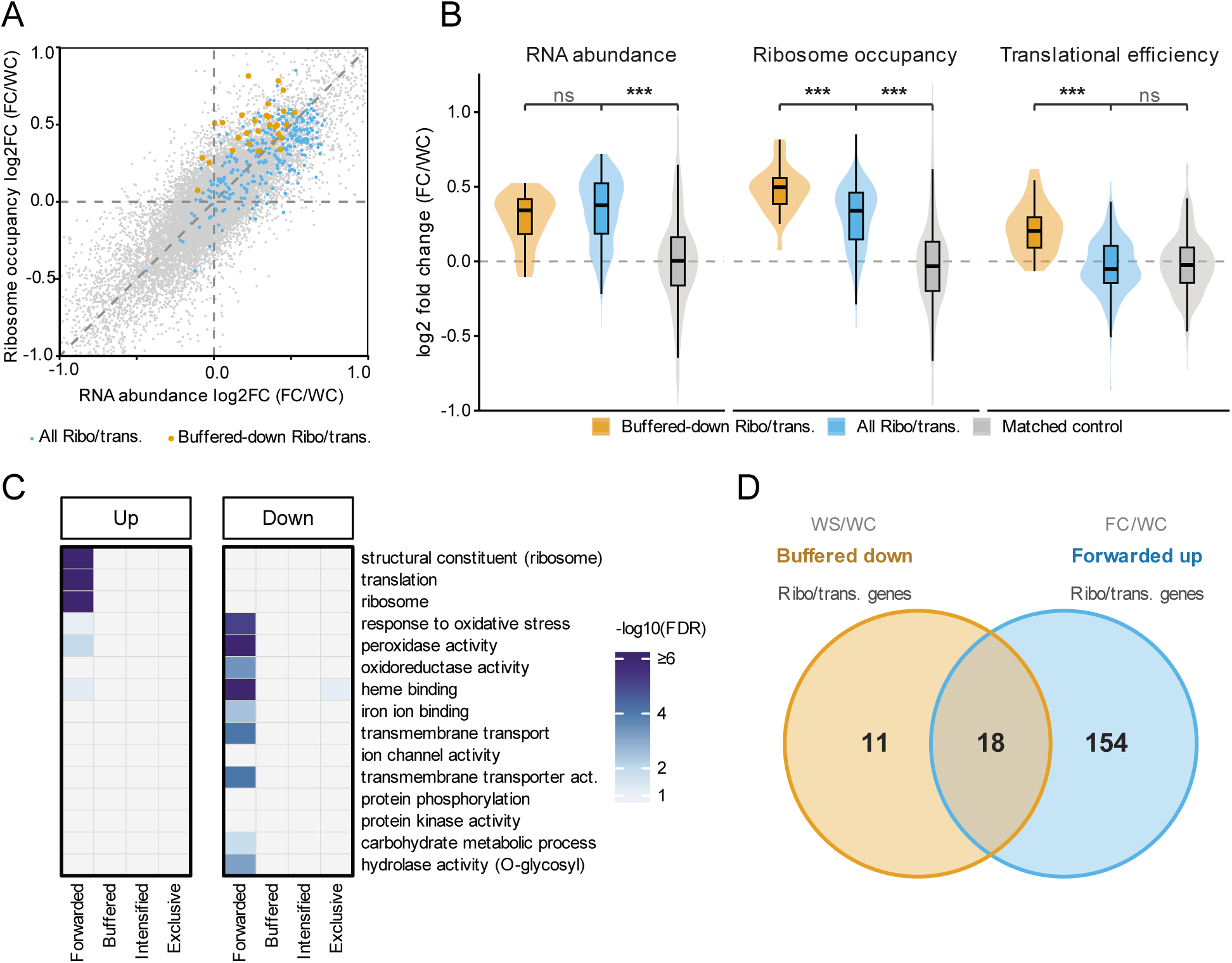
Basal transcriptional and translational differences between *fer* and WT tomato roots. **(A)** RNA and ribosome occupancy changes in the FC/WC comparison, highlighting ribosome/translation GO-annotated genes and the buffered-down subset from the WS/WC comparison (Figure 2). **(B)** RNA abundance, ribosome occupancy and translational efficiency fold changes for the buffered-down ribosome/translation subset, all ribosome/translation GO-annotated genes and matched control genes. **(C)** Gene Ontology enrichment of genes in each regulatory class. Up/down subsets for Forwarded, Buffered and Intensified classes are defined by RNA log_2_FC direction; for the Exclusive class, by Ribo log_2_FC direction. **(D)** Overlap between WS/WC buffered-down and FC/WC forwarded-up ribosome/translation genes. Asterisks in (B): *** *P <* 0.001; Wilcoxon rank-sum tests for Buffered-down versus all ribosome/translation genes and paired Wilcoxon tests for all ribosome/translation genes versus matched controls.

The enrichment of ribosome/translation terms in the FC/WC Forwarded-up class suggested that FER loss shifts the basal state of a translation-related program. To determine whether this basal program overlapped with the WT salt-buffered module, we intersected the WS/WC Buffered-down and FC/WC Forwarded-up ribosome/translation subsets (Figure 3D). Eighteen of 29 Buffered-down ribosome/translation genes (62%) were also Forwarded-up in FC/WC, alongside 154 additional FC/WC Forwarded-up ribosome/translation genes outside the Buffered-down set. This overlap suggests a possible association between FER and the normal WT salt-responsive buffering of the ribosome/translation module.

We next examined the FC/WC gene-level behaviour of both all ribosome/translation GO-annotated genes and the WS/WC Buffered-down ribosome/translation genes. In the FC/WC RNA versus ribosome occupancy scatter plot, 246 of 284 all ribosome/translation GO-annotated genes were located in the upper-right quadrant, consistent with the enrichment of ribosome/translation terms in the FC/WC Forwarded-up class (Figure 3A). The WS/WC Buffered-down ribosome/translation genes showed a more distinct pattern: 26 of 29 were located above the diagonal, indicating that their ribosome-occupancy increases exceeded their RNA increases and suggesting a positive TE shift in this subset. To quantify these layer-specific patterns in FC/WC, we compared RNA abundance, ribosome occupancy and TE log_2_FC values among the WS/WC Buffered-down ribosome/translation subset, all ribosome/translation GO-annotated genes and ribosome-occupancy-matched control genes (Figure 3B). All ribosome/translation GO-annotated genes showed significantly higher RNA abundance and ribosome occupancy than matched controls (P < 0.001), whereas TE did not differ significantly, consistent with coordinated RNA/Ribo up-regulation without a broad TE shift. In contrast, the Buffered-down subset showed significantly higher ribosome occupancy and TE than all ribosome/translation GO-annotated genes (*P <* 0.001), while RNA abundance was similarly elevated between the two groups.

### 2.4 Genotype-by-Salt Interaction Reveals FER-Dependent Ion Transport and Ribosome-Related Translation

Pairwise comparisons above characterised the WT salt response and the *fer* basal transcription–translation state, but cannot isolate regulatory programs whose salt responsiveness specifically depends on FER. We therefore first characterised the *fer* salt response (FS/FC) to establish context, then applied genotype-by-salt interaction analysis to identify FER-dependent programs across RNA, ribosome occupancy and TE layers.

Salt treatment of *fer* roots (FS/FC) elicited a substantial transcriptional and translational response: 894 up- and 613 down-regulated genes were identified at the transcript level, and 977 up- and 521 down-regulated genes at the ribosome occupancy level (Supplementary Figure S2E), with 71.3% of induced and 56.2% of repressed genes shared across both layers (Supplementary Figure S2F). The large majority of regulated genes were classified as Forwarded (*n* = 3,646), with smaller Buffered (*n* = 63), Intensified (*n* = 46), and Exclusive (*n* = 60) classes (Supplementary Figure S3C). All four regulatory classes were smaller than their counterparts in the WS/WC comparison. Unlike the WS/WC comparison, where ribosome/translation terms were enriched in the Buffered-down class, the FS/FC comparison did not show a comparable ribosome/translation enrichment in this class; instead, Supplementary Figure S4A highlighted membrane-, redox-, defence-and cell wall-related terms among the major FS/FC regulatory classes (Table S4). This suggested that FER loss alters the organization of salt-responsive programs, particularly at the level of translational regulation. As additional context before the interaction analysis, we examined the direct genotype comparison under salt conditions (FS/WS). This comparison was similarly dominated by Forwarded genes (*n* = 2,385), with markedly smaller Buffered (*n* = 27), Intensified (*n* = 6) and Exclusive (*n* = 16) classes (Supplementary Figure S3D). GO enrichment of FS/WS regulatory classes showed ribosome/translation terms enriched in the Forwarded-up subset, consistent with the elevated ribosomal mRNA baseline in *fer*persisting under salt, while redox-, heme- and iron-binding functions were strongly enriched in the Forwarded-down class, and ion-transport terms appeared in the Buffered-up subset (Supplementary Figure S4B; Table S4). To directly identify regulatory programs whose salt responsiveness differed between WT and *fer*, we applied gene set enrichment analysis (GSEA) to the genotype-by-salt interaction term.

GSEA was applied across all three comparisons (WS/WC, FC/WC and genotype-by-salt interaction) and regulatory layers; the full bubble-plot overview is shown in Supplementary Figures S5 and S6, and complete GSEA results are provided in Table S5. Focusing on the interaction term, 15 GO terms showed significant genotype-by-salt effects (FDR < 0.05; Figure 4A). Leading-edge overlap analysis highlighted two major functional modules among these interaction-associated terms (Figure 4B). The ribosome/translation-related module, including structural constituent of ribosome, ribosome, translation and intracellular structure, showed significant negative interaction signals in the Ribo and TE layers. Because the interaction contrast was defined as the *fer* salt response minus the WT salt response, these negative NES values indicate that Ribo and TE responses for this module were attenuated in *fer*. The ion-transport module, including cation transport, metal ion transport and metal ion transmembrane transporter activity, showed positive interaction signals mainly in the RNA and Ribo layers, indicating stronger salt-induced RNA abundance and ribosome-occupancy responses of ion-transport genes in *fer* than in WT. These interaction-associated GO terms only partially overlapped with significant terms from the WS/WC and FC/WC comparisons, indicating that the interaction analysis captured FER-dependent salt-responsive programs beyond the pairwise effects (Figure 4C).

**Figure 4:**
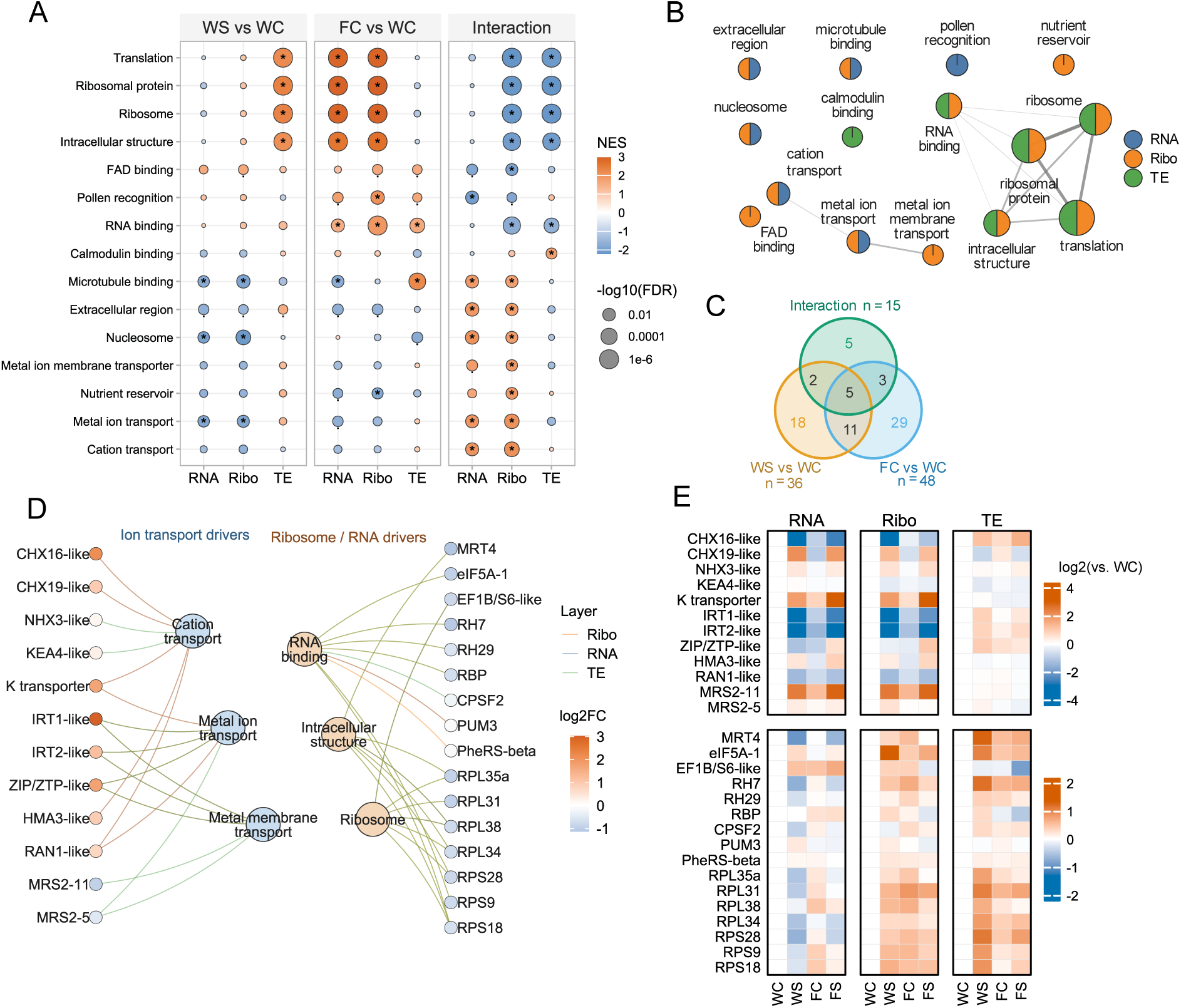
Genotype-dependent salt-response programs in WT and *fer* tomato roots. **(A)** GSEA of interaction-associated GO terms across RNA abundance, ribosome occupancy and translational efficiency. Dot color indicates normalized enrichment score (NES). **(B)** Leading-edge overlap network of significant interaction-associated GO terms. Edges connect GO terms sharing leading-edge genes; node sectors indicate significant RNA, RPF or TE layers. **(C)** Overlap of significant GO terms across the WS/WC, FC/WC and genotype-by-salt interaction comparisons. **(D)** Representative candidate genes from the two major interaction-associated modules in (B). Edge colors indicate RNA, RPF or TE leading-edge connections; gene-node colors indicate RPF interaction log_2_FC for ion-transport genes and TE interaction log_2_FC for ribosome/RNA-related genes. **(E)** RNA abundance, ribosome occupancy and translational efficiency log_2_ ratios relative to WC for ion transport (top) and ribosome/RNA-related (bottom) candidate genes. Symbols in (A): * FDR < 0.05,. FDR < 0.20; significant terms in (B) and (C) were defined by FDR < 0.05.

To move from the two module-level islands in the leading-edge overlap network to their gene-level composition, we visualized representative leading-edge genes from each module (Figure 4D; Table S6). The ion-transport island contained cation/proton exchangers together with metal-, K^+^-and Mg^2+^-transport-related genes, whereas the ribosome/RNA island contained ribosomal proteins and factors associated with translation, ribosome biogenesis/assembly and RNA metabolism. Layer-specific log_2_FC profiles relative to WC further illustrated the behaviour of these representative genes (Figure 4E): the upper ion-transport block showed gene-specific RNA and ribosome-occupancy profiles, with many genes showing stronger salt-associated shifts in *fer* than in WT, whereas the lower ribosome/RNA block showed weaker salt-induced TE increases in *fer* than in WT. Together, these results identify ion transport and ribosome/translation-related processes as prominent FER-dependent components of the tomato root salt response.

### 2.5 Ribosome Occupancy at 5^′^UTRs and uORF-Associated Translation Across Salt and Genotype Conditions

Among the ribosome/RNA-related candidates, eIF5A-1 showed a strong WT salt-induced TE increase that was attenuated in *fer* (Figure 4E). Because eIF5A has been linked to start-codon selection fidelity and suppression of upstream 5^′^UTR translation in other eukaryotes [41], we next examined whether FER-dependent translational changes were accompanied by altered ribosome allocation within 5^′^ leaders and uORF-associated translation. We computed the log_2_ change in relative 5^′^UTR ribosome occupancy, defined as the length-normalized 5^′^UTR-to-CDS ribosome density ratio, for each pairwise comparison. All four comparisons showed significant deviation from zero (one-sample Wilcoxon tests with BH correction, *P <* 0.001; Figure 5A). Under WT salt stress (WS/WC), relative 5^′^UTR ribosome occupancy was significantly reduced. The FS/FC comparison showed a less negative shift than WS/WC (*P* = 1.2 *×* 10^−82^), indicating that the salt-associated reduction in relative 5^′^UTR ribosome occupancy was attenuated in *fer*. This direction is consistent with reports that eIF5A loss increases ribosome occupancy within 5^′^UTRs in other eukaryotes.

**Figure 5:**
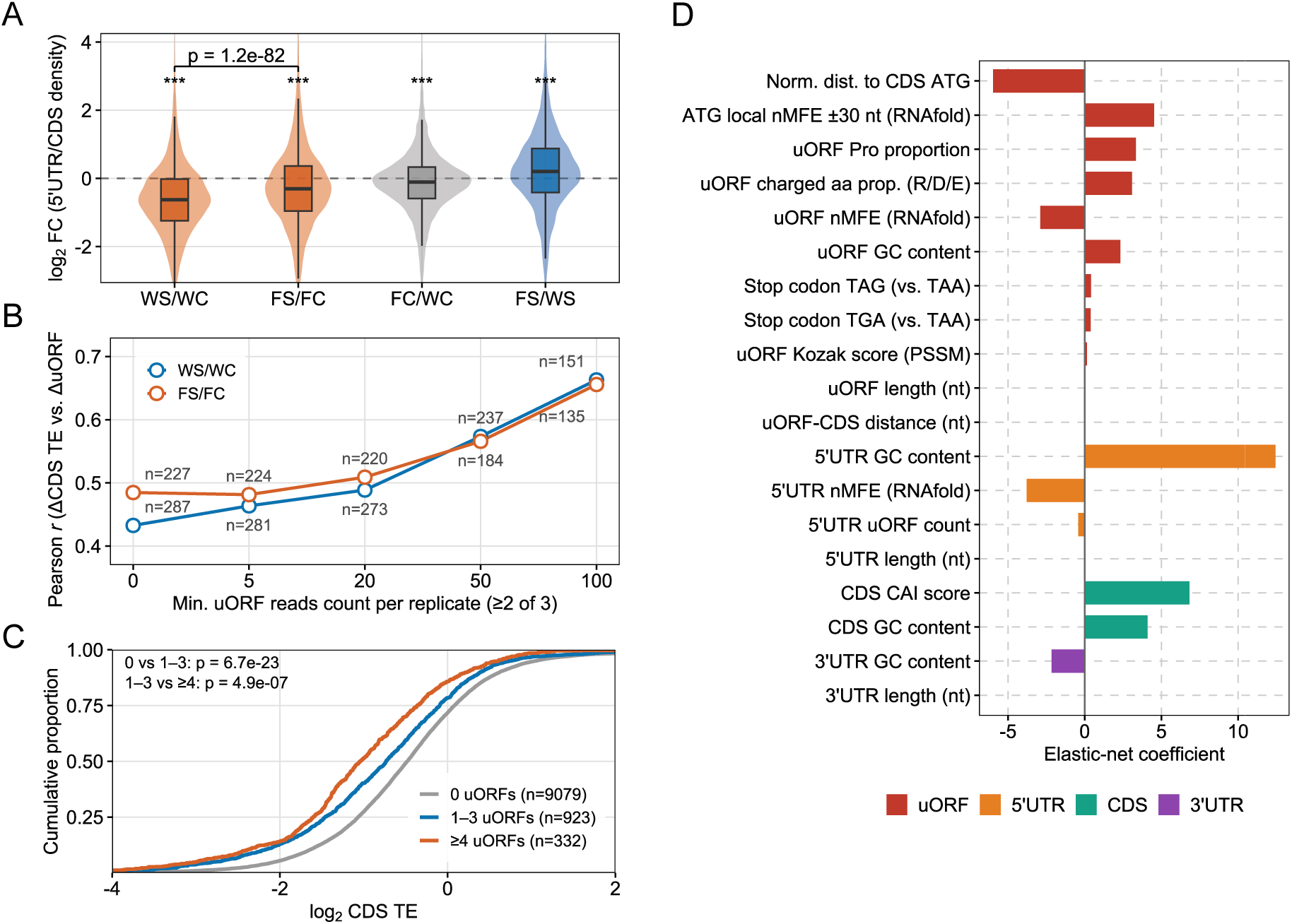
Ribosome occupancy of 5′UTRs and uORF-associated translation in tomato roots. **(A)** Changes in relative 5′UTR ribosome occupancy across salt and genotype comparisons. **(B)** Coupling between CDS and aggregate uORF translational efficiency changes, measured by Pearson correlation, in genes with RPF-supported translated uORFs in both compared conditions. **(C)** Cumulative distributions of CDS translational efficiency among genes grouped by the number of RPF-supported uORFs detected by RiboBA. **(D)** Sequence features distinguishing translated from non-translated uORFs, ranked by elastic-net logistic regression coefficients. Statistics: *** *P <* 0.001 in (A), one-sample Wilcoxon tests with BH correction; bracket in (A), two-sided Wilcoxon rank-sum test; *P* values in (C), pairwise Wilcoxon rank-sum tests with BH correction.

To further dissect the regulatory pattern underlying these 5^′^UTR ribosome shifts, we examined whether aggregate uORF translation changes were competitive or coordinated with downstream main CDS translation. Across genes with RPF-supported translated uORFs in both compared conditions (Table S7), ΔCDS TE and aggregate ΔuORF TE were positively correlated in both WS/WC and FS/FC. Intriguingly, these correlations increased with higher minimum uORF read-count thresholds (Figure 5B), indicating that more robustly detected uORF translation signals showed stronger coupling to downstream CDS TE dynamics. This threshold-dependent reinforcement implies that, at the level of aggregate TE shifts during salt treatment, highly translated uORFs do not predominantly exert a conventional repressive or inhibitory effect on downstream CDS translation; instead, their translational efficiencies are coordinately modulated alongside the main CDS.

While these dynamic ΔTE shifts reveal coordinated uORF–CDS regulation during salt stress, we next asked whether RPF-supported uORF number was associated with downstream CDS TE. After matching for CDS ribosome RPKM, genes with 1–3 RPF-supported uORFs detected by RiboBA had significantly lower CDS TE than genes with no detected uORFs (*P* = 6.7 *×* 10^−23^), and genes with *≥* 4 uORFs showed a further reduction relative to the 1–3 uORF group (*P* = 4.9 *×* 10^−7^; Figure 5C). Thus, RPF-supported uORF number was inversely associated with downstream CDS TE across condition-specific matched sets. Given this association, we next asked which sequence and transcript features distinguish translated from non-translated uORFs. Univariate comparisons revealed broad differences in uORF-, UTR- and CDS-level properties, including uORF length, RNAfold-derived folding metrics, GC content, amino acid composition, distance to the CDS ATG and 5^′^UTR architecture (Supplementary Figure S7). Because these features are interrelated, we used elastic-net logistic regression to rank their joint contributions to distinguishing translated from non-translated uORFs (Figure 5D). The model distinguished translated from non-translated uORFs with high predictive performance (cross-validated AUC = 0.931). The strongest positive predictors were 5^′^UTR GC content and CDS CAI score, followed by local ATG-window nMFE, CDS GC content, uORF proline proportion, uORF charged amino acid proportion and uORF GC content. Negative predictors included normalised distance to the CDS ATG, 5^′^UTR nMFE, uORF nMFE and 3^′^UTR GC content. Together, these results indicate that the distinction between translated and non-translated uORFs is associated with both uORF-intrinsic sequence properties and broader transcript context, with RNAfold-derived folding features contributing in a region-specific manner.

## 3 Discussion

Plant stress responses cannot be inferred from transcript abundance alone, because translational regulation provides an additional regulatory layer [13]. Stress Ribo-seq studies in Arabidopsis hypoxia [42], maize drought [43], rice salinity [15] and potato drought/heat [16] have shown that RNA abundance and ribosome loading are often only partially coupled. Meanwhile, FER has emerged as a receptor kinase linking salt responses with cell-wall integrity, Ca^2+^ signalling and osmotic-stress sensing [26, 44]. This raises the question of whether receptor-mediated signalling contributes to translational remodelling under salt stress. By integrating RNA-seq and Ribo-seq in wild-type and *fer* tomato roots, this study resolved FER-dependent salt responses across transcript abundance, ribosome occupancy and TE layers. The results show that FER loss shifts ribosome/translation-associated genes toward elevated basal transcript abundance and ribosome occupancy, attenuates their salt-induced TE response, and is associated with altered leader-to-CDS ribosome allocation under salt stress.

### 3.1 The WT salt response is transcript-dominant but selectively buffers ribosome related genes

In WT roots, the salt response was transcript-dominant, with Forwarded genes forming the largest regulatory class. The main translational exception was the ribosome/translation-enriched Buffered-down subset: these genes showed salt-induced RNA decreases but concurrent increases in TE, attenuating the expected reduction in ribosome occupancy. Notably, these buffered genes encode components of the protein-synthesis machinery itself. A recent large-scale Ribo-seq/RNA-seq study in mammals showed that translationally buffered genes are enriched for ribosomal proteins and RNA-binding proteins [45]. Our data extend this buffering principle to a plant abiotic-stress setting, identifying ribosome/translation-associated genes as a salt-responsive Buffered-down module in WT tomato roots.

### 3.2 FER deficiency raises basal ribosomal gene expression but reduces salt-induced translational responsiveness

Loss of FER reshaped the basal transcriptional and translational state before salt treatment. FC/WC was dominated by Forwarded regulation, reflecting coordinated RNA and ribosome-occupanc changes without broad TE remodelling. Ribosome/translation-associated genes were coordinately elevated in *fer*, whereas stress-, redox- and transport-associated functions were reduced, indicating a shifted basal expression state before salt treatment.

Notably, 18 of 29 WS/WC Buffered-down ribosome/translation genes were already Forwarded-up in FC/WC, suggesting that part of the WT salt-buffered module is constitutively elevated in *fer* with reduced dynamic range. The genotype-by-salt interaction analysis supported this model: ribosome/translation terms showed attenuated Ribo and TE responses in *fer*, whereas ion-transport terms showed stronger RNA and Ribo responses. These stronger ion-transport responses should not be interpreted as improved tolerance [26, 27], but are more consistent with compensatory or dysregulated stress responses. FER-dependent effects were also evident in ribosome allocation within 5^′^ leaders: the length-normalised 5^′^UTR-to-CDS ribosome density ratio shifted downward under WT salt stress but less so in *fer* (Figure 5A), indicating attenuated redistribution of ribosomes away from leaders in the mutant. Among the ribosome/RNA candidates, *eIF5A-1* provides a potential link to this leader-associated change because its WT salt-induced TE increase was attenuated in *fer*, consistent with reports that eIF5A depletion causes ribosome accumulation within 5^′^UTRs and impairs start-codon fidelity in other eukaryotes [41], although whether FER signalling directly controls eIF5A-1 translational efficiency and whether eIF5A-1 in turn mediates the altered leader-to-CDS ribosome allocation observed in *fer* remain to be established.

### 3.3 Coordinated rather than competitive uORF–CDS translational dynamics during the salt response

The uORF analyses revealed two patterns. First, genes carrying a larger number of RiboBA-detected translated uORF loci showed lower CDS TE even after controlling for CDS ribosome abundance (Figure 5C), consistent with the established view that uORFs can reduce downstream main-ORF translation through impaired scanning or ribosome stalling [30, 35]. Second, aggregate ΔuORF TE and ΔCDS TE were positively correlated across genes with translated uORFs in both compared conditions, and this correlation increased under stricter minimum read-count thresholds (Figure 5B). Similar positive associations between uORF TE and CDS TE have been observed in vertebrate Ribo-seq analyses despite the overall repressive effect of uORFs on CDS translation [46]. In our salt-response data, this positive coupling suggests that the dominant dynamic effect for many genes is likely altered ribosome recruitment across 5^′^ leaders and CDSs rather than direct uORF-mediated repression of downstream CDS translation. One possible explanation is that some translated uORFs are evolutionarily tolerated because their encoded nascent peptides carry regulatory potential. In plants, conserved peptide uORFs can regulate downstream translation through peptide-dependent ribosome stalling or metabolite-responsive control, including stress-responsive CPuORFs and thermospermine-responsive *SAC51* /*SACL* uORFs [32, 33].

### 3.4 Sequence and transcript features distinguish translated uORFs

Single-feature comparisons are useful for screening candidate determinants of translation, but they are limited because sequence and transcript properties are often interrelated [43]. The multivariate feature model therefore provides a more appropriate framework for evaluating the combined determinants of uORF translation (Figure 5D).

Several predictors aligned with established models of eukaryotic 5^′^ leader translation. Stronger Kozak context scores were positively associated with uORF translation, consistent with the well-established role of start-codon context in initiation-site selection [47]. The positive coefficient of local ATG-window nMFE suggests that weaker predicted secondary structure around the uORF start codon favours uORF initiation. Normalised distance to the CDS was negative, indicating that translated uORFs tended to be closer to the main ORF; this is consistent with previous work showing that uORF–CDS spacing modulates downstream translational output [30, 46]. More broadly, sequence features such as GC content and length have been linked to translational output in plant Ribo-seq studies [43].

Beyond these, the model revealed three aspects not straightforwardly predicted from existing frameworks. First, RNAfold-derived nMFE features showed region-specific effects: while weaker predicted structure near the uORF start codon favoured translation, 5^′^UTR and uORF-body nMFE were negative predictors, indicating that translated uORFs were associated with more stable predicted structures outside the immediate start-codon window. Thus, RNA structure showed region-specific effects rather than a uniform inhibitory relationship. Second, 5^′^UTR GC content and CDS CAI were the two largest positive predictors overall, suggesting that transcript-level sequence context contributes to whether a uORF is translated [43, 48]. This was particularly informative for CDS CAI, which showed little univariate separation between translated and non-translated uORFs (BH-adjusted *P* = 0.207) but became a strong positive predictor after joint modelling, underscoring the need for a multivariate framework. Third, translated uORFs were enriched for proline and charged amino-acid proportions, pointing to possible uORF-intrinsic peptide features. These amino-acid patterns are consistent with plant conserved peptide uORFs that can mediate ribosome stalling or regulatory interactions through their encoded nascent peptides [30, 33]. Together, these results indicate that RPF-supported uORF translation in tomato reflects a combination of host-transcript context, start-codon environment, region-specific RNA structure and uORF peptide composition.

## 4 Conclusions

This study provides a matched transcriptome–translatome dissection of FER-dependent salt responses in tomato roots. In WT roots, salt stress was dominated by transcript-driven remodelling, but a ribosome/translation-associated gene module was selectively buffered at the translational level, indicating that translational control refines rather than replaces the transcriptional response. FER deficiency shifted this module toward a higher constitutive baseline and attenuated its salt-induced TE response, while strengthening ion-transport responses in a pattern consistent with a dysregulated stress state. Leader-associated analyses further showed that WT salt stress reduced relative 5^′^UTR ribosome occupancy, an effect attenuated in *fer*, and that aggregate uORF and CDS TE changes were positively coupled during salt treatment. Feature modelling indicated that translated uORFs are distinguished by transcript context, start-codon environment, region-specific RNA structure and uORF peptide composition. Together, these results support a model in which FER contributes to tomato salt adaptation by maintaining the dynamic range of selective translation, and identify FER signalling, *eIF5A-1* and leader-associated ribosome allocation as candidate mechanistic links for future study.

## 5 Materials and Methods

### 5.1 Plant Materials and Salt Treatment

Wild-type tomato (*Solanum lycopersicum* cv. Condine Red) was obtained from the Tomato Genetics Resource Center at the University of California, Davis. The CRISPR/Cas9-generated *fer* mutant line in the cv. Condine Red background was generated as described previously [24].

Seedlings were grown in a greenhouse under controlled conditions with a 16 h light/8 h dark photoperiod at 30^◦^C during the light period and 25^◦^C during the dark period. Plants were maintained in 1/4-strength Hoagland’s medium before treatment. At 3–4 weeks after sowing, seedlings were divided into control and salt-treatment groups. For the salt treatment, plants were subjected to a progressive salt treatment culminating in 150 mM NaCl; control plants were maintained in 1/4-strength Hoagland’s medium. Root tissues were harvested after 5 days of salt treatment from each group, with three biological replicates per treatment, and used for parallel RNA-seq and ribosome profiling analyses.

### 5.2 RNA-seq and Ribosome Profiling Library Preparation

Total RNA was extracted from tomato roots using the RNAprep Pure Plant Kit (DP419; Tiangen Biotech, Shanghai, China) according to the manufacturer’s instructions. RNA-seq libraries were then constructed following a standard mRNA-seq library preparation workflow and sequenced in paired-end mode with a read length of 2 *×* 150 nt.

Ribosome profiling libraries were prepared using a modified Ribo-seq workflow. Briefly, clarified lysates were digested with RNase I and DNase I to generate ribosome-protected fragments (RPFs), and ribosome-associated fractions were recovered using a MicroSpin S-400 column. RPFs were then extracted, followed by rRNA depletion and RNA cleanup. The purified fragments were subjected to 5^′^ phosphorylation with T4 polynucleotide kinase, ligated sequentially to 3^′^ and 5^′^ adaptors, reverse transcribed, and PCR amplified for 12 cycles. The final libraries were size selected by PAGE, and fragments of approximately 140 bp were recovered for sequencing.

### 5.3 Processing, Alignment and Quantification of RNA-seq and Ribo-seq Data

Raw Ribo-seq reads were processed with fastp (v0.20.1) [49] using paired-end merging and adapter correction. Merged reads of 25–40 nt were retained. Reads derived from rRNA were removed by aligning to tomato rRNA sequences with Bowtie (v1.3.1) [50] using the parameter -v 0. Non-rRNA reads were mapped to the tomato SL4.0 genome with ITAG4.0 annotation using STAR (v2.7.10b) [51] with –alignEndsType EndToEnd –outFilterMismatchNmax 1 –quantMode TranscriptomeSAM. Mapping efficiency was calculated as genome-mapped reads divided by input reads after rRNA removal.

Ribo-seq gene-level ribosome occupancy was quantified from CDS-overlapping reads using featureCounts in the Rsubread package (v2.22.1) [52]. RNA-seq reads were aligned to the same reference genome using STAR in two-pass mode, and exon-level gene counts were obtained with featureCounts. Multi-mapping and multi-overlap reads were excluded. For sample-level visualization, raw count matrices were normalized by the TMM method and transformed to logCPM using edgeR (v4.6.3) [53].

### 5.4 Reproducibility, PCA and Ribo-seq Quality Control

For reproducibility analysis, genes with counts *≥* 10 in at least three samples were retained. Sample reproducibility was assessed using squared Pearson correlation coefficients calculated from TMM-normalized logCPM values. Principal component analysis was performed on the same filtered and normalized RNA-seq and Ribo-seq gene-level matrices.

Ribo-seq P-site calibration and quality-control analyses were performed with riboWaltz (v2.0) [54]. Transcriptome-aligned BAM files were filtered to retain uniquely mapped reads and converted to riboWaltz-compatible input using the ITAG4.0 annotation. P-site offsets were inferred automatically and used to summarize read-length distribution, CDS reading-frame preference, read distribution across 5^′^UTR, CDS and 3^′^UTR regions, and metagene profiles around annotated start and stop codons.

### 5.5 Differential Expression, Translational Efficiency and Regulatory Classification

Differential transcript-abundance and ribosome-occupancy analyses were performed with DESeq2 (v1.48.2) [55]. Gene-level TE was defined as normalized ribosome occupancy divided by normalized RNA abundance following the deltaTE framework [14]. Translational regulatory classes were assigned as follows: Forwarded genes showed significant transcript-abundance and ribosome-occupancy changes without significant TE changes; Buffered genes showed transcript-abundance and TE changes in opposite directions; Intensified genes showed transcript-abundance and TE changes in the same direction; and Exclusive genes showed significant ribosome-occupancy and TE changes without significant transcript-abundance changes. For directional summaries, Forwarded, Buffered and Intensified classes were split by RNA log_2_FC direction, whereas Exclusive genes were split by ribosome-occupancy log_2_FC direction.

### 5.6 GO Enrichment and Gene Set Enrichment Analysis

GO over-representation analysis was performed with clusterProfiler [56] using ITAG4.0-derived GO annotations. The background was defined as the set of genes entering the corresponding deltaTE analysis.

Gene set enrichment analysis (GSEA) was performed with fgsea [57]. For genotype-by-salt interaction analysis, genes were ranked by interaction scores calculated as (FS*/*FC) *−* (WS*/*WC) separately for RNA abundance, ribosome occupancy and TE. GO terms containing 10–500 genes were retained, and fgsea was run with nPermSimple = 10000. Leading-edge genes from terms passing the GSEA FDR threshold were used to construct the GO-term overlap network and candidate-gene views.

### 5.7 uORF Identification and Leader-associated Translation Analysis

Sequence-defined uORFs and RPF-supported uORF translation events were identified using RiboBA [38] from ITAG4.0 transcript annotation and raw Ribo-seq data. RPF-supported uORF events were summarized by identical transcript-coordinate intervals, defined by transcript, strand, uORF start and uORF end. For translated-uORF feature modelling, translated uORFs were defined as uORFs with the same transcript-coordinate interval detected in at least two biological replicates in any of the four conditions.

For relative 5^′^UTR ribosome occupancy, RiboBA-derived P-site counts were summed separately over 5^′^UTR and CDS regions and divided by region length to calculate P-site density. The length-normalized ratio was defined as 5^′^UTR P-site density divided by CDS P-site density. Gene-level condition means were calculated across three biological replicates, and log_2_ fold changes of this ratio were computed for WS/WC, FS/FC, FC/WC and FS/WS.

To examine uORF–CDS TE coupling, genes carrying RPF-supported translated uORFs in both conditions of a comparison were retained. Aggregate uORF P-site signal was calculated by summing P-site counts across uORF intervals within each gene, normalized to CPM, and divided by RNA CPM to obtain aggregate uORF TE. CDS TE was calculated analogously from CDS P-site CPM and RNA CPM. Pearson correlations between Δaggregate uORF TE and ΔCDS TE were calculated across a series of minimum raw uORF P-site thresholds.

For CDS TE cumulative distribution analysis, genes were grouped in each condition by the number of RPF-supported uORFs detected by RiboBA in at least one replicate: 0, 1–3 or *≥* 4 uORFs. To control for CDS ribosome abundance, genes in the 0 and 1–3 groups were matched to the *≥* 4 group by binned mean CDS ribosome RPKM. Matched condition-specific gene sets were then combined across the four conditions.

### 5.8 Sequence-feature Analysis and Elastic-net Modelling of Translated uORFs

Sequence and transcript features were extracted for translated and non-translated ATG-uORFs from transcript sequences using in-house scripts. Non-translated uORFs were obtained by scanning 5^′^UTRs of matched control genes for ATG-initiated ORFs fully contained within the 5^′^UTR, excluding intervals overlapping translated uORFs. Features included uORF length, uORF–CDS distance, normalized distance to the CDS ATG, uORF and UTR GC content, RNAfold-derived [58] normalized minimum free energy (nMFE) for the uORF, 5^′^UTR and local ATG window, uORF start-codon Kozak score, uORF amino-acid composition, 5^′^UTR uORF number, stop codon type and CDS CAI score. The Kozak score was calculated as a position-specific scoring matrix score from the nucleotide context around annotated CDS start codons and then applied to uORF start-codon contexts [47]. CAI was calculated according to the codon adaptation index framework [48].

Feature contributions were modelled using elastic-net binomial logistic regression implemented in glmnet [59]. Predictors were median-imputed where necessary and standardized during model fitting. The model used *α* = 0.5, with *λ* selected by 10-fold cross-validation using AUC as the optimization metric. Predictive performance was evaluated by cross-validation and an internal held-out test set, and non-zero coefficients at the minimum cross-validated lambda were used to rank features distinguishing translated from non-translated uORFs.

### 5.9 Statistical Analysis

Unless otherwise stated, statistical tests were two-sided. Fresh-weight differences between treatments and genotypes were assessed by Wilcoxon rank-sum tests. For transcript-abundance and ribosome-occupancy DEG summaries, differentially regulated genes were defined by adjusted *p <* 0.05 and *|* log_2_ FC*| ≥* 1. For deltaTE regulatory classification, transcript-abundance, ribosome-occupancy and TE components were considered significant at adjusted *p <* 0.05.

For GO over-representation analysis, Benjamini–Hochberg-adjusted *P* values were used, with significance thresholds as indicated in the corresponding figures. For GSEA, FDR < 0.05 was used as the significance threshold; FDR < 0.20 is shown as a suggestive level in Figure 4A.

Gene-set comparisons used Wilcoxon rank-sum tests for unpaired groups or paired Wilcoxon signed-rank tests for matched-control comparisons. For Figure 5A, one-sample Wilcoxon signed-rank tests against zero were Benjamini–Hochberg corrected, whereas the WS/WC versus FS/FC comparison used a Wilcoxon rank-sum test. CDS TE distributions grouped by uORF count and uORF sequence-feature comparisons were tested by pairwise Wilcoxon rank-sum tests with Benjamini–Hochberg correction; stop-codon type was evaluated by chi-squared test.

## Supplementary Materials

The following supporting information can be downloaded at: Supplementary Materials, Figure S1: Sequencing quality assessment of RNA-seq and Ribo-seq libraries. **(A)** Pairwise *R*^2^ correlation heatmaps of Ribo-seq and RNA-seq libraries across 12 samples from WC, WS, FC and FS conditions, with three biological replicates per condition. **(B)** Reading frame distribution of CDS P-sites across the four experimental groups. **(C)** Distribution of P-sites across 5^′^UTR, CDS and 3^′^UTR regions for Ribo-seq libraries and RNA-seq reads; Figure S2: RNA abundance and ribosome occupancy changes across salt and genotype comparisons. **(A, C, E)** Numbers of up- and down-regulated genes at transcriptional and translational levels in the WS/WC, FC/WC and FS/FC comparisons, respectively. **(B, D, F)** Overlap between transcriptionally and translationally regulated genes for the corresponding up- and down-regulated subsets. Differential genes were defined by adjusted *P <* 0.05 and *|* log_2_ FC*| ≥* 1; Figure S3: Relationship between RNA abundance and translational efficiency changes across salt and genotype comparisons. **(A–D)** RNA abundance and translational efficiency changes in the WS/WC, FC/WC, FS/FC and FS/WS comparisons, respectively. Genes were colored according to regulatory classes defined by RNA, RPF and TE changes; Figure S4: Gene Ontology enrichment of regulatory classes in *fer* salt and genotype comparisons. **(A)** Gene Ontology enrichment of Forwarded, Buffered, Intensified and Exclusive classes in the FS/FC comparison. **(B)** Gene Ontology enrichment in the FS/WS comparison. Colored cells indicate enriched terms (FDR < 0.10); color intensity represents *−* log_10_(FDR); Figure S5: GSEA overview of genotype, salt and interaction effects across RNA, RPF and TE layers (part 1). GSEA bubble plots showing all GO terms with FDR < 0.05 in at least one layer of at least one comparison, across the WS/WC, FC/WC and genotype-by-salt interaction effects. Bubble color indicates NES and bubble size indicates *−* log_10_(FDR). Asterisks indicate FDR < 0.05; Figure S6: GSEA overview of genotype, salt and interaction effects across RNA, RPF and TE layers (part 2). Continuation of Figure S5. Bubble color indicates NES and bubble size indicates *−* log_10_(FDR). Asterisks indicate FDR < 0.05; Figure S7: Sequence and transcript features of translated and non-translated uORFs. Distributions of uORF, UTR and CDS sequence features compared between translated and non-translated uORFs. Translated uORFs were defined by detection of the same transcript-coordinate interval in at least two replicates in any condition. BH-adjusted *P* values are shown for Wilcoxon rank-sum tests, except for uORF stop codon type, which was tested by chi-square test; Table S1: Alignment statistics for Ribo-seq libraries; Table S2: Alignment statistics for RNA-seq libraries; Table S3: Significant differential transcript abundance, ribosome occupancy, translational efficiency and WS/WC regulatory class assignments; Table S4: Gene Ontology enrichment results for regulatory classes across salt and genotype comparisons; Table S5: Gene set enrichment analysis results for salt, genotype and genotype-by-salt interaction effects; Table S6: Gene-level source tables for ribosome/translation-associated buffered modules and interaction candidate genes; Table S7: Transcript-coordinate translated uORFs detected in at least two replicates in any condition.

## Author Contributions

J.B. and Y.F. conceived and designed the study. Y.F. performed plant cultivation, salt treatment, tissue collection and RNA-seq and Ribo-seq library preparation. J.B. performed bioinformatics and statistical analyses and generated all figures. R.Y. and J.Y. supervised the study. J.B. drafted the manuscript. All authors have read and agreed to the published version of the manuscript.

## Institutional Review Board Statement

Not applicable.

## Informed Consent Statement

Not applicable.

## Data Availability Statement

RNA-seq and Ribo-seq data for wild-type and *fer* tomato roots under salt stress are available in ArrayExpress under E-MTAB-17154 and E-MTAB-17159. Figure-ready data and plotting code are available in Zenodo under DOI: https://doi.org/10.5281/zenodo.20563675.

## Acknowledgments

During the preparation of this manuscript, the authors used OpenAI Codex (GPT-5.3) and Anthropic Claude Sonnet 4.6 to assist with data-processing pipeline organization, statistical analysis and figure generation. The authors reviewed, edited and tested the output and take full responsibility for the content of this publication.

## Abbreviations

The following abbreviations are used in this manuscript: 
AUC: area under the curve
CAI: codon adaptation index
CDS: coding sequence
DEG: differentially expressed gene
FC: *fer* control
FDR: false discovery rate
FER: FERONIA
FS: *fer* salt
GO: Gene Ontology
GSEA: gene set enrichment analysis
logCPM: log counts per million
nMFE: normalised minimum free energy
NES: normalised enrichment score
PCA: principal component analysis
Ribo-seq: ribosome sequencing
RNA-seq: RNA sequencing
RPF: ribosome-protected fragment
TE: translational efficiency
TMM: trimmed mean of M-values
uORF: upstream open reading frame
WC: WT control
WS: WT salt (150 mM NaCl)
WT: wild-type

