## Supplementary Information for "FERONIA-Dependent Translational Buffering of Ribosome-Associated Genes in Salt-Stressed Tomato Roots"

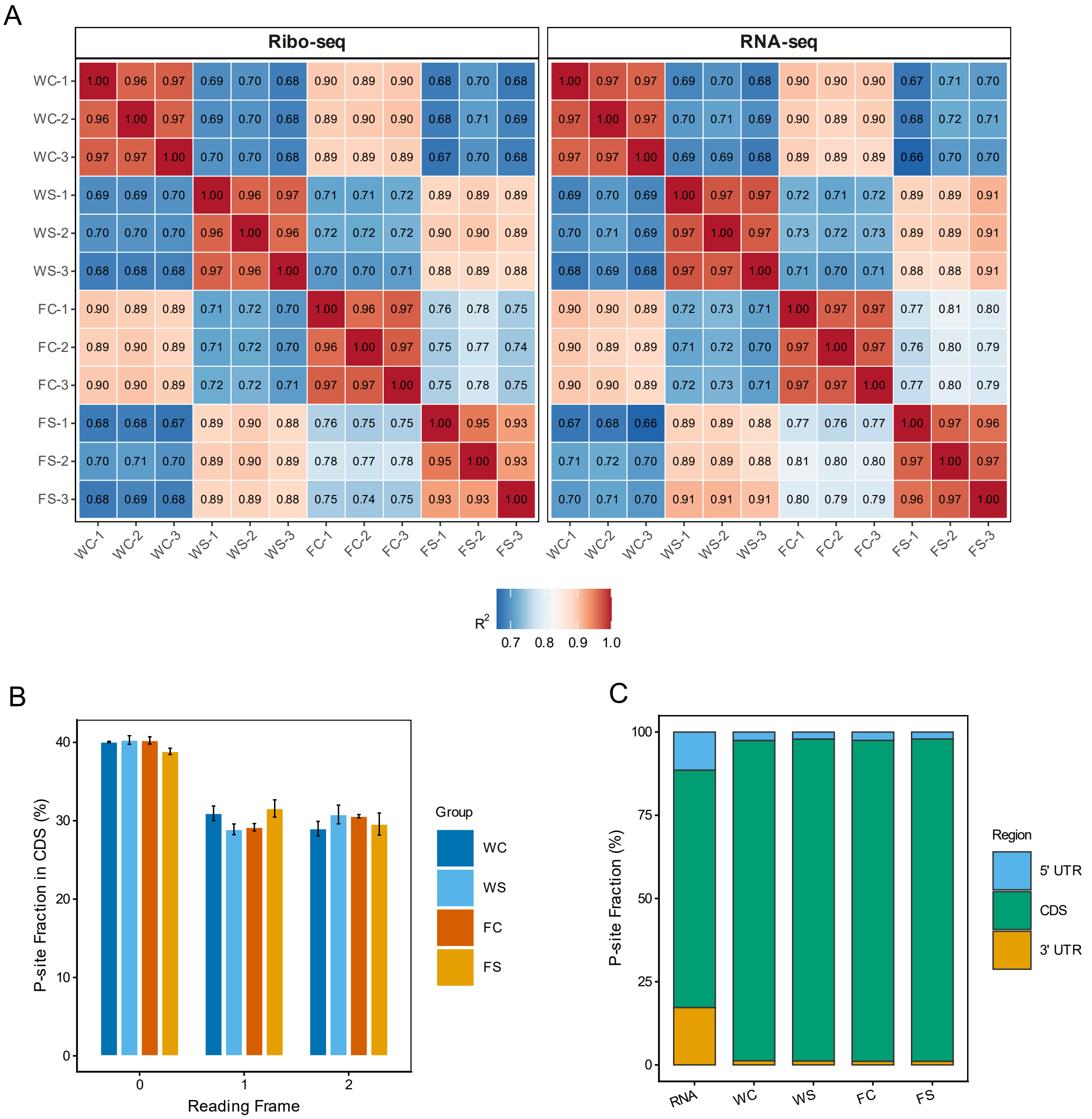


**Supplementary Figure S1. Sequencing quality assessment of RNA-seq and Ribo-seq libraries. (A)** Pairwise $R^2$ correlation heatmaps of Ribo-seq and RNA-seq libraries across 12 samples from WC, WS, FC and FS conditions, with three biological replicates per condition. **(B)** Reading frame distribution of CDS P-sites across the four experimental groups. **(C)** Distribution of P-sites across 5'UTR, CDS and 3'UTR regions for Ribo-seq libraries and RNA-seq reads.

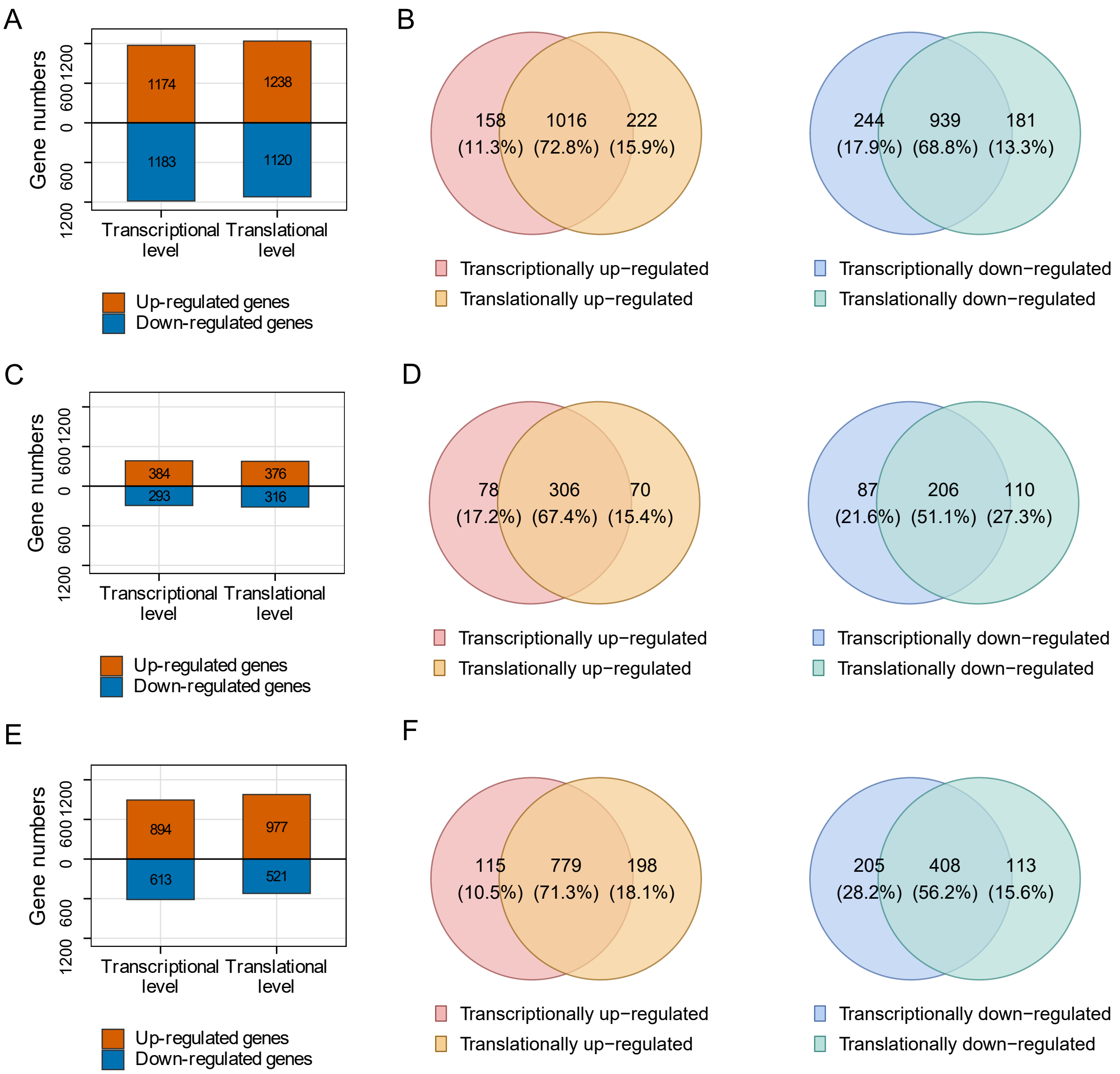


**Supplementary Figure S2. RNA abundance and ribosome occupancy changes across salt and genotype comparisons. (A, C, E)** Numbers of up- and down-regulated genes at transcriptional and translational levels in the WS/WC, FC/WC and FS/FC comparisons, respectively. **(B, D, F)** Overlap between transcriptionally and translationally regulated genes for the corresponding up- and down-regulated subsets. Differential genes were defined by adjusted $P < 0.05$ and $|\log_2 \mathrm{FC}| \geq 1$.

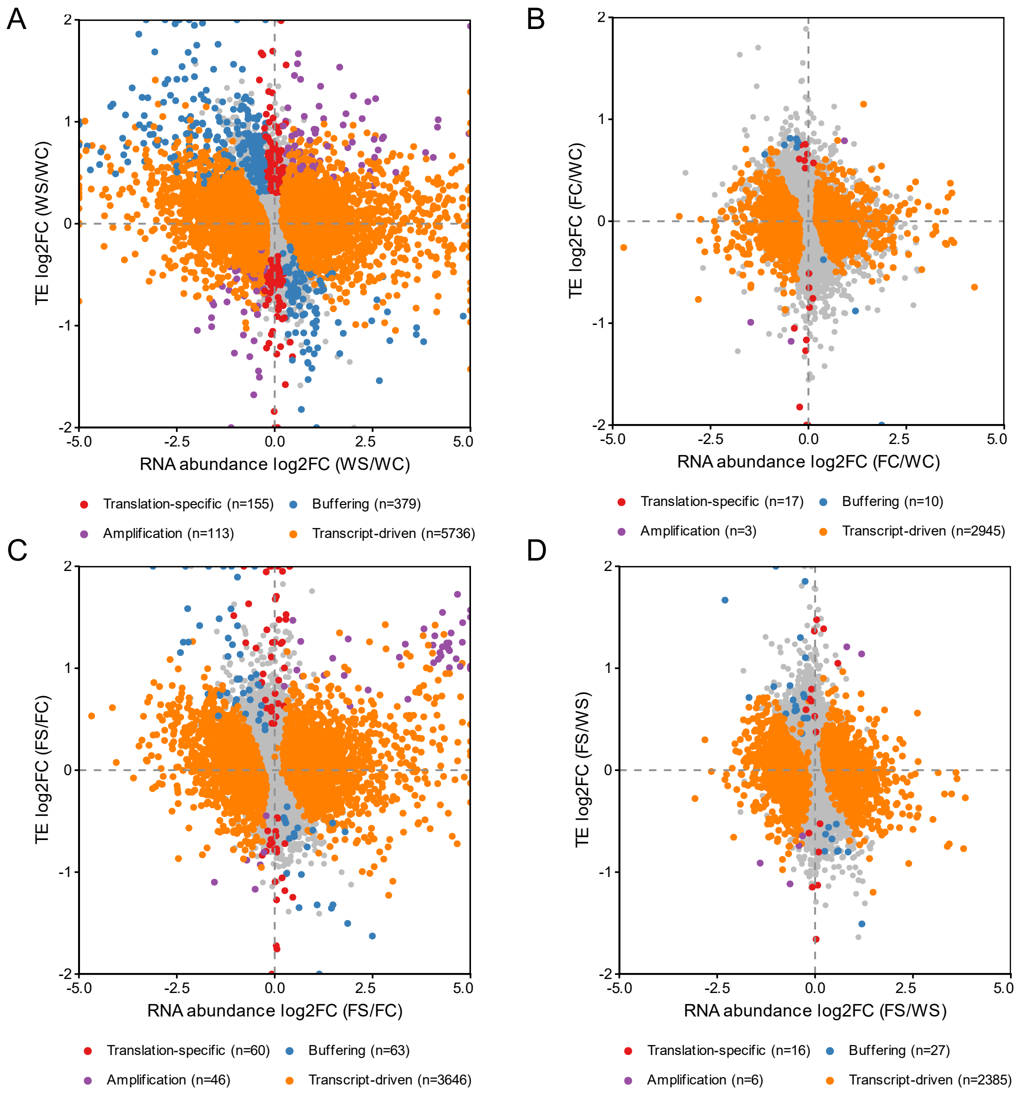


**Supplementary Figure S3. Relationship between RNA abundance and translational efficiency changes across salt and genotype comparisons. (A–D)** RNA abundance and translational efficiency changes in the WS/WC, FC/WC, FS/FC and FS/WS comparisons, respectively. Genes were colored according to regulatory classes defined by RNA, RPF and TE changes. Class labels correspond to Forwarded (Transcript-driven), Buffered (Buffering), Intensified (Amplification) and Exclusive (Translation-specific) as defined in Figure 2.

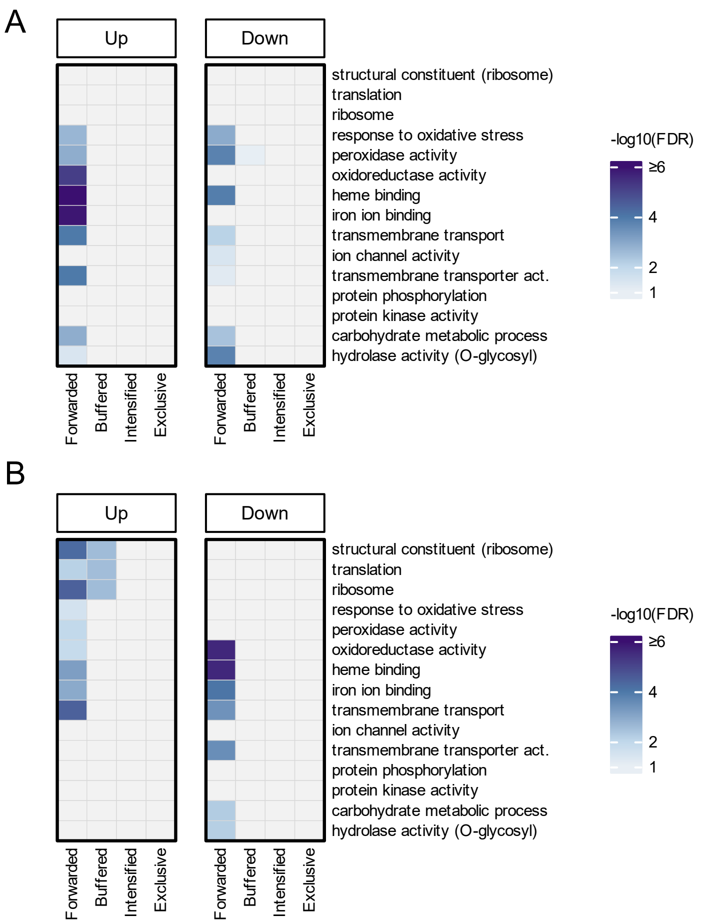


**Supplementary Figure S4. Gene Ontology enrichment of regulatory classes in *fer* salt and genotype comparisons. (A)** Gene Ontology enrichment of Forwarded, Buffered, Intensified and Exclusive classes in the FS/FC comparison. **(B)** Gene Ontology enrichment in the FS/WS comparison. For Forwarded, Buffered and Intensified classes, up- and down-regulated subsets are defined by RNA $\log_2$FC direction; for the Exclusive class, by Ribo $\log_2$FC direction. Colored cells indicate enriched terms (FDR $< 0.10$); color intensity represents $-\log_{10}(\mathrm{FDR})$.

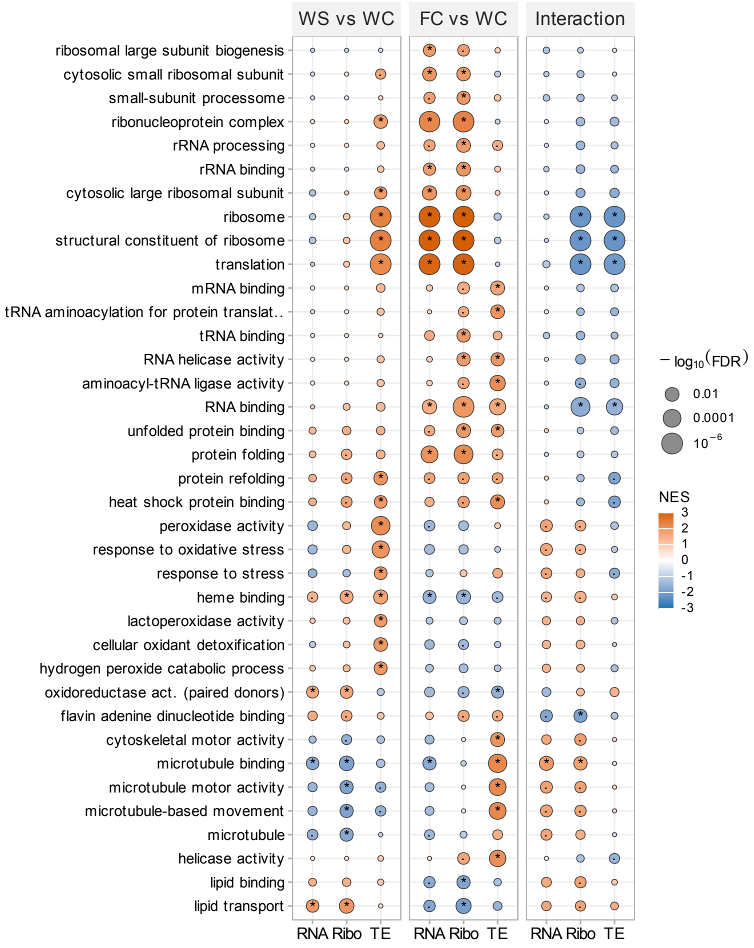


**Supplementary Figure S5. GSEA overview of genotype, salt and interaction effects across RNA, RPF and TE layers (part 1).** GSEA bubble plots showing all GO terms with FDR $< 0.05$ in at least one layer of at least one comparison, across the WS/WC, FC/WC and genotype-by-salt interaction effects in RNA, RPF and TE layers. Bubble color indicates normalized enrichment score (NES), and bubble size indicates $-\log_{10}(\mathrm{FDR})$. Asterisks indicate FDR $< 0.05$.

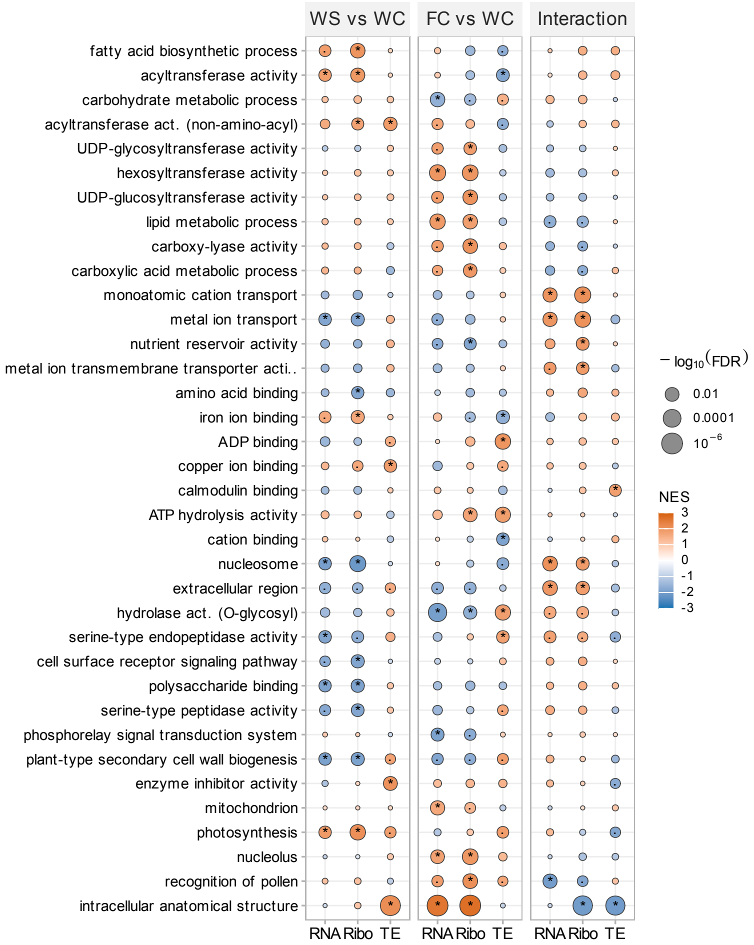


**Supplementary Figure S6. GSEA overview of genotype, salt and interaction effects across RNA, RPF and TE layers (part 2).** Continuation of Supplementary Figure S5. Bubble color indicates normalized enrichment score (NES), and bubble size indicates $-\log_{10}(\mathrm{FDR})$. Asterisks indicate FDR $< 0.05$.

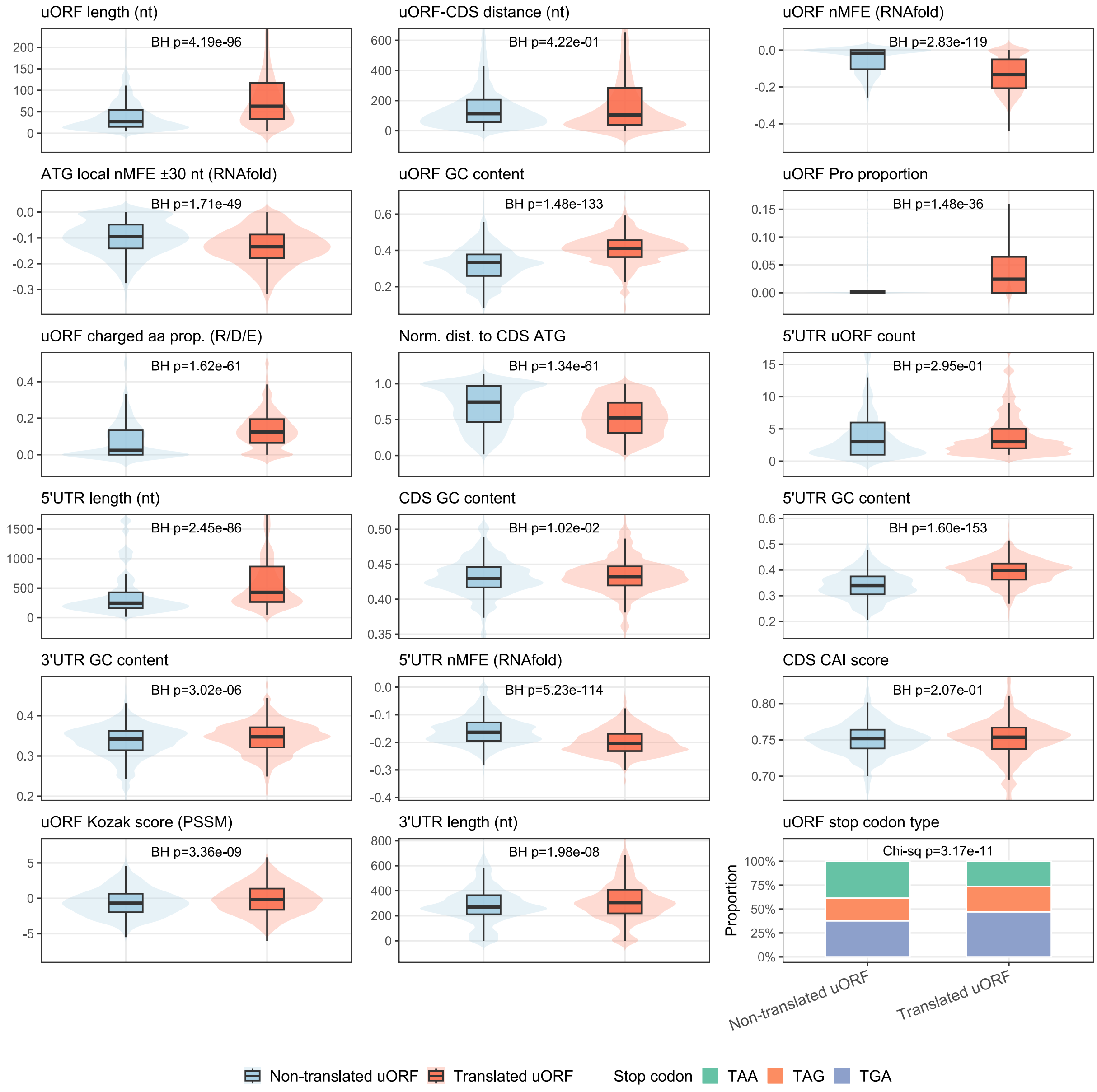


**Supplementary Figure S7. Sequence and transcript features of translated and non-translated uORFs.** Distributions of uORF, UTR and CDS sequence features compared between translated and non-translated uORFs. Translated uORFs were defined by detection of the same transcript-coordinate interval in at least two replicates in any condition. BH-adjusted $P$ values are shown for Wilcoxon rank-sum tests, except for uORF stop codon type, which was tested by chi-square test.
